# Genome-wide association analysis of 350,000 Caucasians from the UK Biobank identifies novel loci for asthma, hay fever and eczema

**DOI:** 10.1101/195933

**Authors:** Åsa Johansson, Mathias Rask-Andersen, Torgny Karlsson, Weronica E. Ek

**Affiliations:** Department of Immunology, Genetics and Pathology, Science for Life Laboratory, Uppsala University, Uppsala. Sweden

## Abstract

Even though heritability estimates suggest that the risk of asthma, hay fever and eczema is largely due to genetic factors, previous studies have not explained a large part of the genetics behind these diseases. In this GWA study, we include 346,545 Caucasians from the UK Biobank to identify novel loci for asthma, hay fever and eczema. We further investigate if associated lead SNPs have a significantly larger effect for one disease compared to the other diseases, to highlight possible disease specific effects.

We identified 141 loci, of which 41 are novel, to be associated (P≤3×10^−8^) with asthma, hay fever or eczema, analysed separately or as disease phenotypes that includes the presence of different combinations of these diseases. The largest number of loci were associated with the combined phenotype (asthma/hay fever/eczema). However, as many as 20 loci had a significantly larger effect on hay fever/eczema-only compared to their effects on asthma, while 26 loci exhibited larger effects on asthma compared with their effects on hay fever/eczema. At four of the novel loci, *TNFRSF8, MYRF, TSPAN8*, and *BHMG1*, the lead SNPs were in LD (> 0.8) with potentially casual missense variants.

Our study shows that a large amount of the genetic contribution is shared between the diseases. Nonetheless, a number of SNPs have a significantly larger effect on one of the phenotypes suggesting that part of the genetic contribution is more phenotype specific. Identified loci and probable causal genes may in the future be used as targets for treatments of asthma, hay fever and eczema.

## Introduction

Asthma, hay fever and eczema are common complex immunological diseases affecting many people worldwide (1). The prevalence for these diseases vary among populations and have an underlying architecture that include both environmental and genetic risk factors (1). Comorbidity between asthma, hay fever and eczema is common, and previous genome-wide association (GWA) studies have, apart from identifying a large number of genetic variants associated with risk of disease (2–8) also found evidence of a genetic overlap between the diseases (6, 9).

Family and twin studies have estimated that the contribution of genetic factors, i.e. the heritability for asthma (1, 10, 11) to be 35-95%, for hay fever (1, 11) to be 33-91%, and for eczema (12) to be as high as 90%. A recent large study estimated the SNP-based heritability, the heritability that can be attributed the genetic variation captured by SNPs in a GWA study, to be 15% for asthma, 22% for hay fever, and 9% eczema (6). The same study performed a GWA study that included the first release of UK Biobank (N=138,354) analysing asthma, hay fever and eczema as a combined phenotype and identified 99 significantly associated loci (6). Many of the identified target genes were predicted to influence the function of immune cells, and only six loci were identified to have disease specific effects (6). Many previous GWA studies for asthma, hay fever and eczema have been conducted in different cohorts that were subsequently meta-analysed with the purpose of increasing statistical power (2, 4, 6–8, 13).

The aim with this study was to explain a larger part of the genetic background of self-reported asthma, hay fever, and eczema as well as identify possible novel disease specific effects. We investigated the genetic background of self-reported asthma, hay fever, and eczema combined to a single phenotype, similar as to Ferreira *et al* (6), but in a more homogenous population, as we included both the first and the second release of UK Biobank (N=346,545), compared to Ferreira (6) that only included the first UK Biobank release (N=138,354) as part of a large meta-analysis. We also analysed each disease phenotype independently in larger groups than previous studies conducted in UK Biobank. We further had a larger power to investigated if associated lead SNPs had a significantly larger effect for one disease phenotype compared to the other phenotypes, to highlight possible disease specific effects. Associated SNPs were functionally annotated to assess likely causal mechanisms.

Although the phenotypes in the UK Biobank are self-reported, the questions are well defined and identical for all participants.

## Results

### Association analysis

The UK Biobank includes 502,682 participants, of which 443,068 are Caucasians. The disease prevalence in the Caucasian participants were 11.7% for asthma and 23.2% for hay fever and/or eczema (combined). As many as 45.8% of the asthmatic participants had reported having hay fever and/or eczema, and 23.0% of the hay fever and/or eczema participants had reported being diagnosed with asthma (Figure 2). We conducted the GWA study using 346,545 unrelated Caucasians (Table 1), who passed the quality control (QC) for the second UK Biobank genetic data release, and had no ambiguities with regards to disease status. After QC, 15,688,218 genetic variants where included in the analyses (see methods). QC and the final number of included participants in respective analyses are summarized in the methods and in Table 1. We did not identify any statistically significant associations located on the X-chromosome.

**Figure 1.**
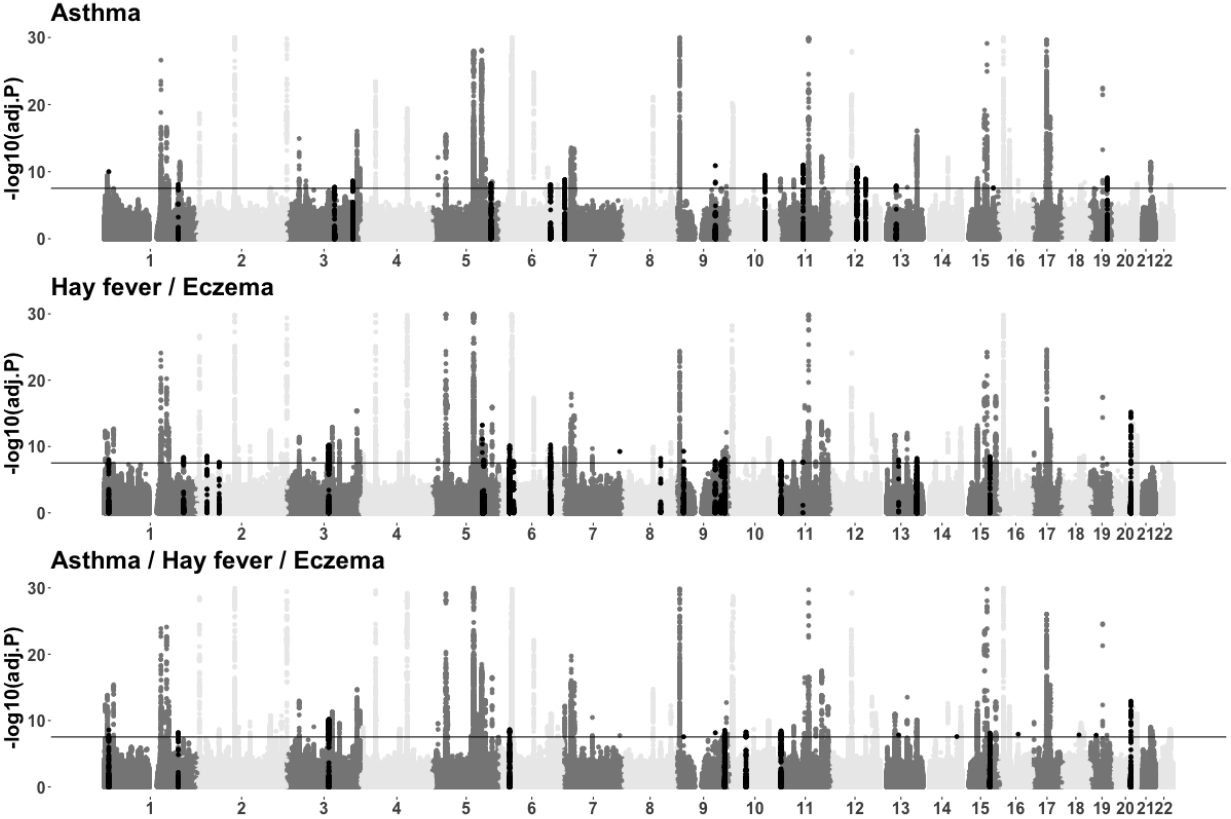
Manhattan plots for asthma, for hay fever and/or eczema, and for asthma and/or hay fever and/or eczema (combined) for autosomal chromosomes. The black horizontal line indicates the genome wide threshold (3×10^−8^). The black regions represent novel loci found in this study.

**Figure 2.**
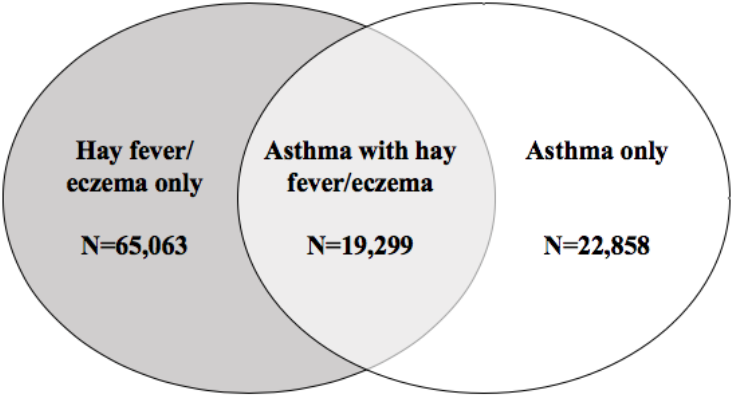
Comorbidity between asthma and hay fever/eczema.

**Table 1.**
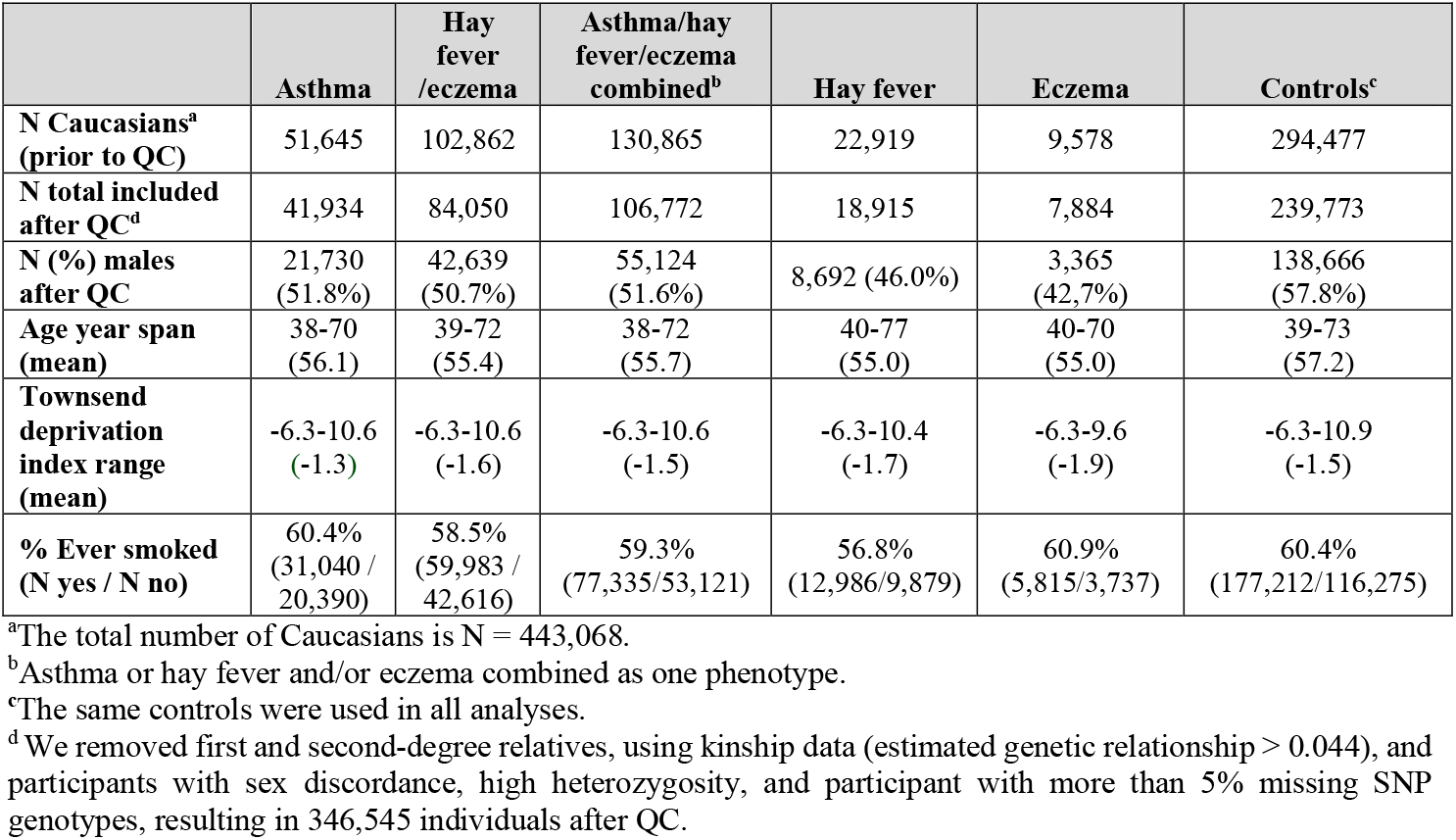
Baseline characteristics of Caucasian participants in UK Biobank.

|  | **Asthma** | **Hay fever /eczema** | **Asthma/hay fever/eczema combined<sup>b</sup>** | **Hay fever** | **Eczema** | **Controls<sup>c</sup>** |
| --- | --- | --- | --- | --- | --- | --- |
| **N Caucasians<sup>a</sup> (prior to QC)** | 51,645 | 102,862 | 130,865 | 22,919 | 9,578 | 294,477 |
| **N total included after QC<sup>d</sup>** | 41,934 | 84,050 | 106,772 | 18,915 | 7,884 | 239,773 |
| **N (%) males after QC** | 21,730 (51.8%) | 42,639 (50.7%) | 55,124 (51.6%) | 8,692 (46.0%) | 3,365 (42.7%) | 138,666 (57.8%) |
| **Age year span (mean)** | 38-70 (56.1) | 39-72 (55.4) | 38-72 (55.7) | 40-77 (55.0) | 40-70 (55.0) | 39-73 (57.2) |
| **Townsend deprivation index range (mean)** | -6.3-10.6 (-1.3) | -6.3-10.6 (-1.6) | -6.3-10.6 (-1.5) | -6.3-10.4 (-1.7) | -6.3-9.6 (-1.9) | -6.3-10.9 (-1.5) |
| **% Ever smoked (N yes / N no)** | 60.4% (31,040 / 20,390) | 58.5% (59,983 / 42,616) | 59.3% (77,335/53,121) | 56.8% (12,986/9,879) | 60.9% (5,815/3,737) | 60.4% (177,212/116,275) |

<sup>a</sup>The total number of Caucasians is N = 443,068.
<sup>b</sup>Asthma or hay fever and/or eczema combined as one phenotype.
<sup>c</sup>The same controls were used in all analyses.
<sup>d</sup>We removed first and second-degree relatives, using kinship data (estimated genetic relationship > 0.044), and participants with sex discordance, high heterozygosity, and participant with more than 5% missing SNP genotypes, resulting in 346,545 individuals after QC.

### GWA study for self-reported asthma

After QC, 41,926 self-reported asthma cases (independent on hay fever/eczema status) and 239,773 controls were included in the GWA analysis. We identified 75 risk loci located > 1 Mb apart and containing at least one significantly associated genetic variant (P≤3×10^−8^ after adjusting for LD-score intercept of 1.065), that were associated with self-reported asthma, of which 15 loci were found to be novel asthma loci not previously identified in a GWA study (Table 2; Manhattan plot, Figure 1; S1 Table and S2 Table; quantile-quantile (QQ) plot, S1 Fig). Using approximate conditional analysis (14), we identified 116 independent significant associations within these 75 loci (S1 Table). The strongest associations for asthma were found within the HLA locus on chromosome 6 (P=2.06×10^−100^), including 14 independent genetic variants. Several genes within this region have previously been reported to be associated with asthma (i.e., *HLA-DQB1, HLA-G* and *HLA-DRB1*) (1, 3, 13). Among the novel asthma loci, some have previously been associated with other similar phenotypes (S1 Table). For example, *SDK1*, previously annotated to the nearby *CARD11* gene, have been reported to be associated with atopic dermatitis (15), but this is the first time that the *SDK1* locus has been identified in a GWA study for asthma. Five of the 15 novel lead SNPs were further replicated in an independent cohort (P<0.05; Table 2). However, most of the SNPs were not possible to investigate in the replication cohort, due to a low number of overlapping SNPs between the cohorts.

**Table 2.**
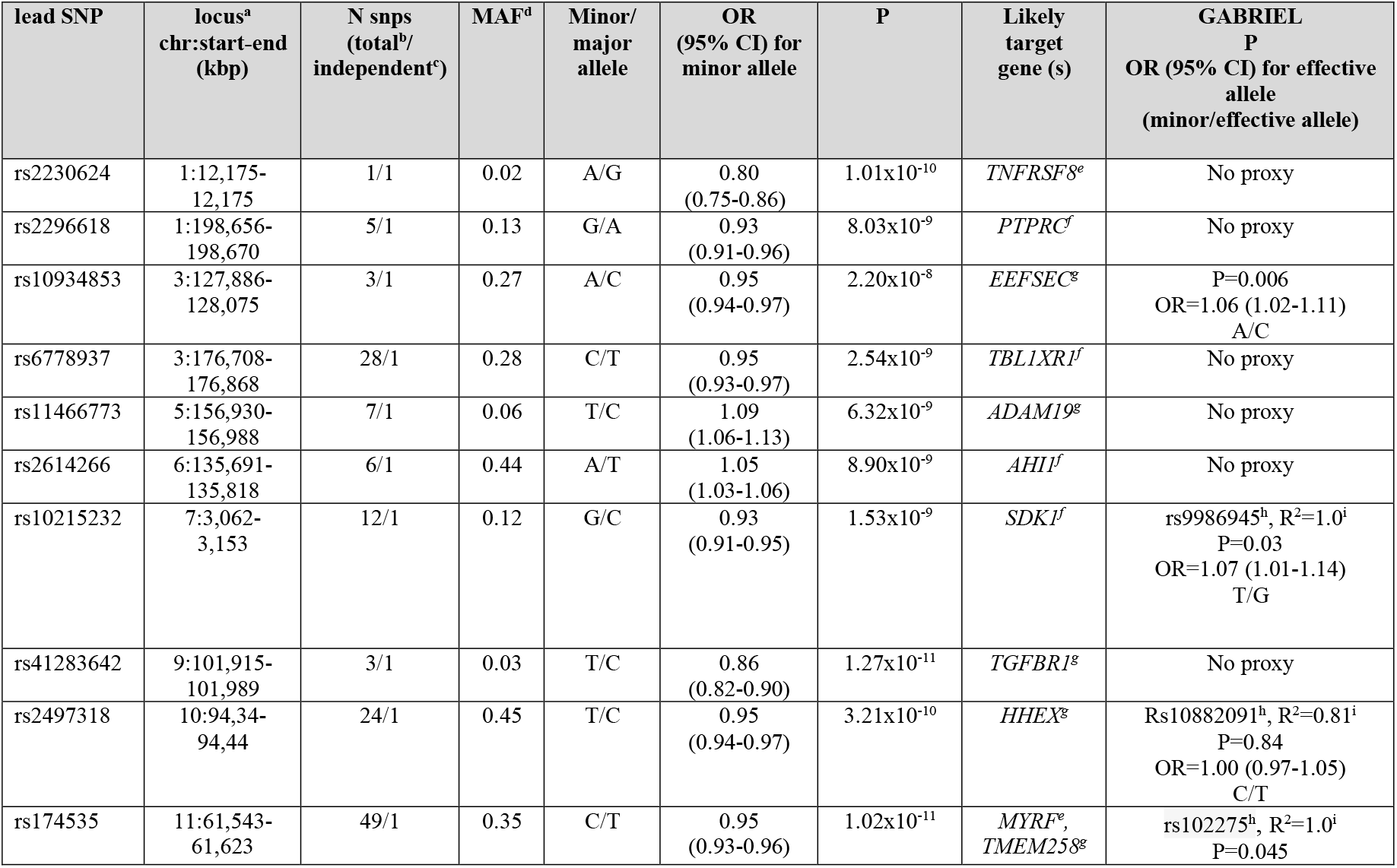
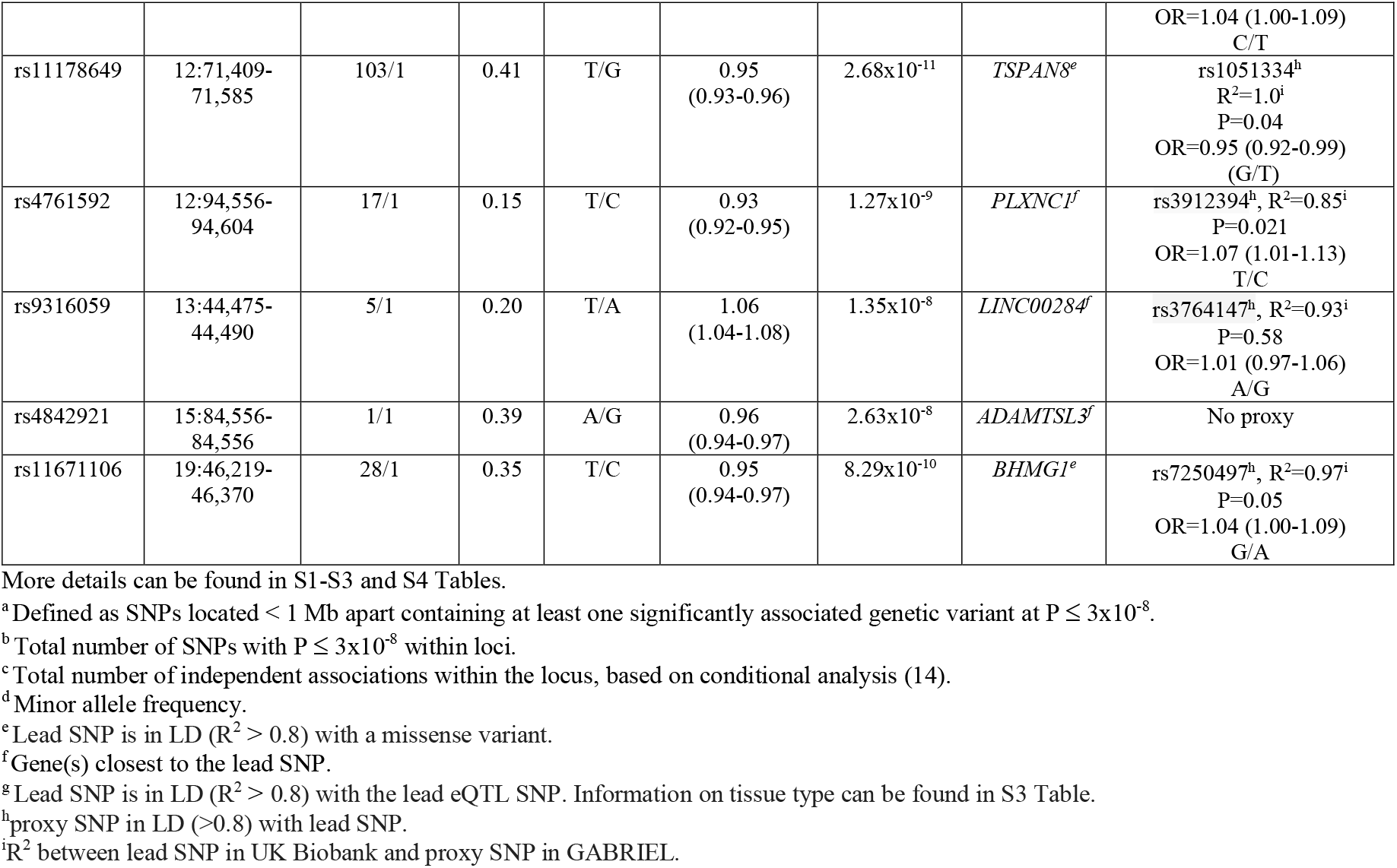
Summary results for the 15 novel loci significantly associated with self-reported asthma in UK Biobank (P ≤ 3×10^−8^) with replication in the GABRIEL cohort.

### Annotation of asthma associated SNPs

Associated SNPs were further functionally annotated to assess likely causal mechanisms (see Methods). Overlap with GTEx eQTLs were found for 15 of the 75 asthma loci. Of these, four eQTLs (*EEFSEC, ADAM19, HHEX* and *TMEM258*) overlapped with the novel loci, where increased expression of *TMEM258* in cell transformed fibroblasts appears to lower the risk for developing asthma (Table 2; S1 Table S1 and S3 Table). In contrast, increased expression of *EEFSEC* in lung tissue seems to increase the risk for asthma (S3 Table). Similarly, increased expression of *ADAM19* in whole blood and *HHEX* in cell transformed fibroblasts appears to increase the risk of asthma (S3 Table). 19 probable causal missense variants could be observed within the 75 significant GWA loci of which four missense variants for the 15 novel loci (S4 Table). The latter are located within *TNFRSF8, MYRF, TSPAN8*, and *BHMG1*. The association at *TNFRSF8* was represented by only one genetic variant, rs2230624 (S2 Fig). This SNP is a missense variant in two transcripts for *TNFRSF8* and causes a cysteine to a tyrosine substitution which was predicted as ‘probably damaging’ by PolyPhen (16) (PolyPhen-score 0.751-0.921) and had a ‘deleterious’ SIFT-score (17) of 0. The lead SNP at the *MYRF* locus, rs174535, is a missense variant in five transcripts for *MYRF*. Rs174535 causes a serine to arginine substitution and was predicted to be ‘probably damaging’ by PolyPhen(16) (PolyPhen-score 0.961-1) and had a ‘deleterious’ SIFT-score (17) of 0.04-0.07. However, rs174535 is also in LD with the most significant eQTL for *TMEM258* in cell transformed fibroblasts. The lead SNP in the *BHMG1* locus, rs11671106, is a missense variant for *BHMG1* and was predicted as ‘probably damaging’ by PolyPhen (16) (PolyPhen-score 0.94) and had a ‘deleterious’ SIFT-score (17) of 0.01. The lead SNP at the *TSPAN8* locus, rs11178649, was in complete LD with rs3763978 (R^2^=1), a missense variant in three transcripts for *TSPAN8*, which causes a glycine to alanine substitution which was predicted as ‘probably damaging’ by PolyPhen (16) (PolyPhen-score 0.989) and had a ‘deleterious’ SIFT-score(17) of 0.03

### GWA study for self-reported hay fever/eczema

After QC, 84,034 self-reported hay fever and/or eczema cases that were combined as a single phenotype were included in the analysis. We identified 109 loci to be associated (P≤3×10^−8^, LD-score intercept =1.079) with self-reported hay fever/eczema, and 22 of these were novel (Table 3; Manhattan plot, Figure 1; S5 Table and S6 Table; QQ-plot, S3 Fig). The strongest association was observed for the lead SNP rs5743604 (P= 7.5×10^−72^) located within *TLR1*. This SNP has previously been associated with allergic disease (6, 21). Using conditional analysis, we identified 154 independent significant associations within these 109 loci (Table 3; S5 Table). Moreover, two of our lead SNPs (rs4845604 and rs9986945, mapped to *RORC* and *SDK1*), observed within previously known loci were in low LD (R^2^≤0.05) with the previously reported genetic variants, indicating that they represent novel variants within or close to known loci (S5 Table). The *UBAC2* locus has previously been reported to be associated with asthma (9), but this is the time that the *UBAC2* locus is reported to be associated with hay fever and/or eczema. We replicated six of the novel lead SNPs in an independent eczema GWA study, the EAGLE consortium (P≤0.05), (Table 3). However, all SNPs that did not replicate in EAGLE were neither significant in UK Biobank when analysing eczema separately (Table 3).

**Table 3.**
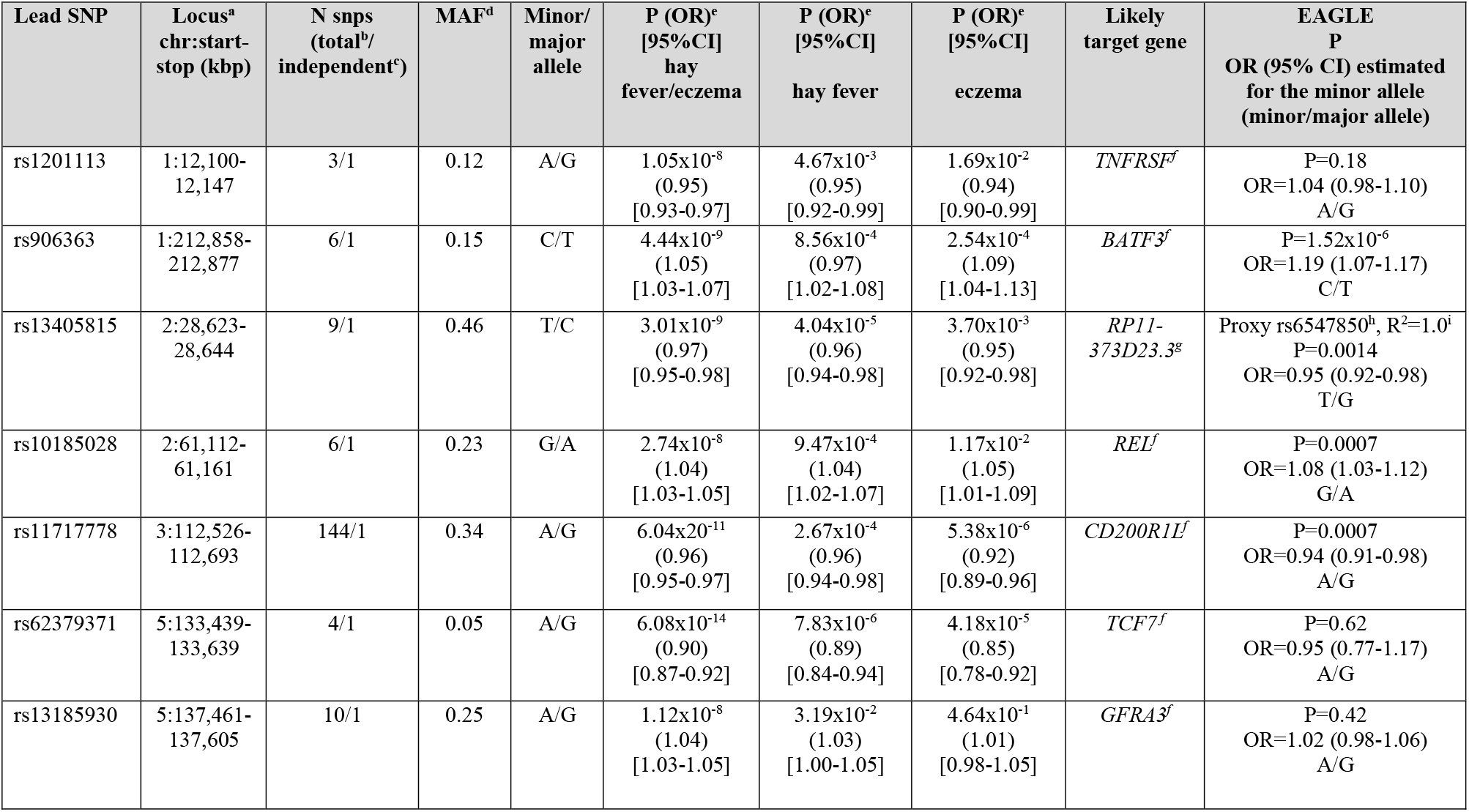
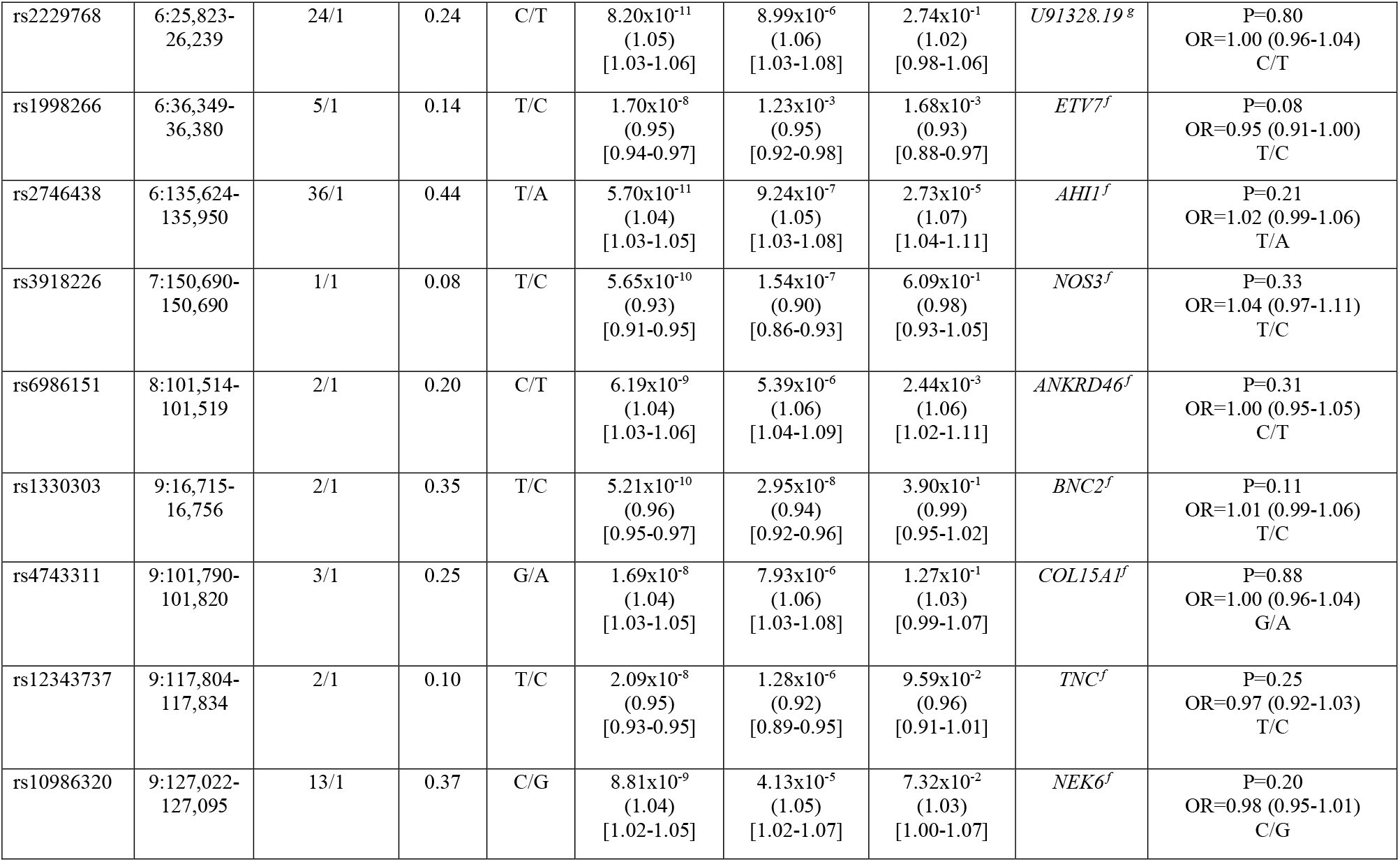
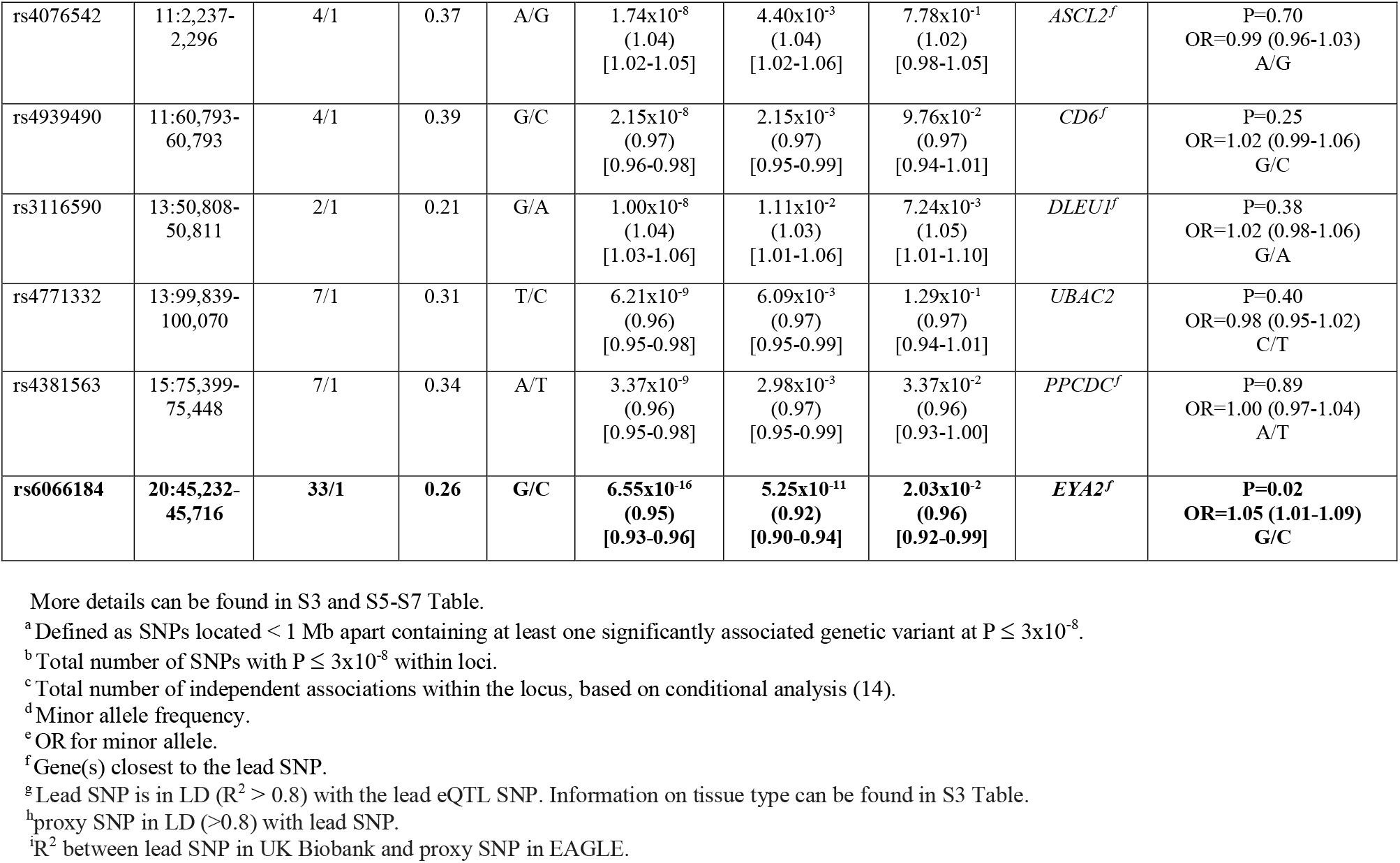
Summary results for the 22 novel loci significantly associated with self-reported hay fever and/or eczema in UK Biobank (P ≤ 3×10^−8^) with replication in the EAGLE cohort

### Annotation of hay fever/eczema SNPs

For eleven of the 109 hay fever/eczema associated loci, the lead SNP was in LD with the lead SNP for GTEx eQTLs (Table 3; S3 Table) and 14 overlapped with possible causal missense variants in genes, including *IL6R, IL7R, IL13* and *SMAD4* (S7 Table).

### GWA studies for hay fever and eczema analysed separately

Hay fever and eczema could not be separated for most of the participants, since they had primarily answered yes or no on whether they had either hay fever or eczema. However, to investigate hay fever and eczema individually, we also analysed hay fever (N hay fever cases = 18,915) and eczema (N eczema cases = 7,884) separately in a smaller subset of UK Biobank participants (Manhattan plot, S4 Fig; QQ-plots, S5 and S6 Fig). A total of 27 and 18 loci were identified for hay fever and eczema, respectively. One novel hay fever and one novel eczema locus, which has not been reported in previous GWA studies and that were not significantly associated in the combined hay fever/eczema analysis, was detected when analysing hay fever and eczema separately (S8-S11 Tables). The lead SNP, rs12920150 (P=1.02×10^−9^), at the hay fever locus is located close to *CBLN1* and the lead SNP, rs2485363 (P=1,20×10^−8^), at the eczema locus is located downstream of *TAGAP*. This novel eczema locus was nominally replicated using the summary statistics from the GWA study on eczema in the EAGLE consortium (P=0.018, OR=1.05 [95% CI 1.02-1.92]). Another locus that was not detected when analysing hay fever/eczema combined was detected when analysing eczema separately. The lead SNP for this locus, rs676387 (P=2.26×10^−10^), is located within *HSD17B1* (S10 and S11 Tables). This region has previously been reported to be associated with allergic disease (6) and overlap with an eQTL for *TUBG2* in skin, where a decreased expression of *TUBG2* seems to lower the risk for eczema (S3 Table).

### GWA study for asthma/hay fever/eczema (combined as a single phenotype)

For the combined analysis of asthma and/or hay fever and/or eczema (N cases=106,752), we identified 110 significant loci (LD-score intercept=1.081), and 16 of these were novel GWA loci that have not been significantly associated with either asthma, hay fever or eczema in previous GWA studies (Table 4; Manhattan plot, Figure 1; S12 and S13 Tables; QQ-plot, S7 Fig). However, 12 of these 16 novel loci were detected when analysing asthma and hay fever/eczema separately, while the remaining four novel loci were only found when analysing asthma, hay fever, and/or eczema together as a single phenotype. Using conditional analysis, we identified 164 independent associations within these 110 loci (Table 4; S12 Table). The most significant SNP, rs72823641 (P=1.14×10^−78^), was located within *IL1RL1* and was also significantly associated with asthma and hay fever/eczema when these phenotypes were analysed separately (P = 4.09×10^−61^ and P = 9.64×10^−64^) (S12 and S13 Tables). This region has previously been associated with allergic diseases (6) (S14 Table). We also identified five lead SNPs for the combined phenotype asthma and/or hay fever and/or eczema within previously known loci, which were found to be in low LD (R^2^≤0.05) with previously reported genetic variants, indicating that they represent novel variants within known loci. These five lead SNPs mapped to *LPP, IL31, LINC00393, CCR7* and *NFATC* (S12 and S14 Tables).

**Table 4.** Summary results for the 16 novel loci significantly associated with self-reported asthma and/or hay fever and/or eczema (combined) in UK Biobank (P ≤ 3×10^−8^).

| Lead SNP | Locus<sup>a</sup><br>chr:start-stop (kbp) | N snps<br>(total<sup>b</sup>/<br>independent<sup>c</sup>) | MAF<sup>d</sup> | Minor/<br>major allele | P (OR<sup>e</sup>)<br>[95%CI]<br>combined | P (OR<sup>e</sup>)<br>[95%CI]<br>asthma | P (OR<sup>e</sup>)<br>[95%CI]<br>hay fever/<br>eczema | Likely<br>target gene |
| --- | --- | --- | --- | --- | --- | --- | --- | --- |
| rs2230624 | 1:12,080-12,175 | 2/2 | 0.02 | A/G | $2.64 \times 10^{-9}$<br>(0.87)<br>[0.84-0.91] | $1.01 \times 10^{-10}$<br>(0.80)<br>[0.75-0.86] | $1.99 \times 10^{-7}$<br>(0.88)<br>[0.84-0.92] | *TNFRSF8<sup>f</sup>* |
| rs7410883 | 1:198,640-198,670 | 5/1 | 0.11 | C/T | $7.00 \times 10^{-9}$<br>(0.95)<br>[0.93-0.97] | $2.26 \times 10^{-8}$<br>(0.93)<br>[0.91-0.95] | $1.78 \times 10^{-6}$<br>(0.96)<br>[0.94-0.97] | *PTPRC<sup>g</sup>* |
| rs9816107 | 3:112,526-112,693 | 160/1 | 0.34 | A/C | $6.80 \times 10^{-11}$<br>(0.96) | $1.51 \times 10^{-6}$<br>(0.96)<br>[0.95-0.97] | $8.39 \times 10^{-11}$<br>(0.96)<br>[0.95-0.97] | *CD200R1L<sup>g</sup>* |
| rs62379371 | 5:133,439-133,639 | 4/1 | 0.05 | A/G | $1.13 \times 10^{-13}$<br>(0.91)<br>[0.89-0.93] | $1.23 \times 10^{-7}$<br>(0.91)<br>[0.88-0.94] | $6.08 \times 10^{-14}$<br>(0.90)<br>[0.87-0.92] | *VDAC1<sup>g</sup>* |
| rs9379828 | 6:26,038-26,184 | 13/1 | 0.37 | G/C | $2.71 \times 10^{-9}$<br>(1.03)<br>[1.02-1.05] | $1.07 \times 10^{-10}$<br>(1.05)<br>[1.04-1.07] | $1.42 \times 10^{-6}$<br>(1.03)<br>[1.02-1.04] | *HIST1H2BD<sup>h</sup>* |
| rs1330303 | 9:16,715-16,715 | 1/1 | 0.35 | T/C | $2.84 \times 10^{-8}$<br>(0.97)<br>[0.96-0.98] | 0.016<br>(0.98)<br>[0.97-1.00] | $5.21 \times 10^{-10}$<br>(0.96)<br>[0.95-0.97] | *BNC2<sup>g</sup>* |
| rs41283642 | 9:101,915-101,915 | 1/1 | 0.03 | T/C | $7.03 \times 10^{-9}$<br>(0.92)<br>[0.89-0.94] | $1.27 \times 10^{-11}$<br>(0.86)<br>[0.82-0.90] | $6.42 \times 10^{-6}$<br>(0.93)<br>[0.90-0.96] | *TGFBR1<sup>g</sup>* |
| rs3758212 | 9:127,002-127,178 | 14/1 | 0.37 | T/C | $2.99 \times 10^{-9}$<br>(1.03)<br>[1.02-1.05] | $1.80 \times 10^{-5}$<br>(1.04)<br>[1.02-1.05] | $2.18 \times 10^{-8}$<br>(1.04)<br>[1.02-1.05] | *NEK6<sup>g</sup>* |
| **rs2505504** | **10:43,728-43,763** | **17/1** | **0.29** | **A/G** | **$5.45 \times 10^{-9}$**<br>**(1.04)**<br>**[1.02-1.05]** | **$1.96 \times 10^{-6}$**<br>**(1.04)**<br>**[1.02-1.06]** | **$4.61 \times 10^{-7}$**<br>**(1.03)**<br>**[1.02-1.05]** | ***RASGEF1A<sup>g</sup>*** |
| rs7114923 | 11:2,237-2,305 | 21/1 | 0.37 | T/C | 3.74x10<sup>-9</sup><br>(1.03)<br>[1.02-1.05] | 1.20x10<sup>-5</sup><br>(1.04)<br>[1.02-1.05] | 2.65x10<sup>-8</sup><br>(1.04)<br>[1.02-1.05] | *ASCL2*<sup>g</sup> |
| rs3116590 | 13:50,808-50,808 | 1/1 | 0.21 | G/A | 1.52x10<sup>-8</sup><br>(1.04)<br>[1.03-1.05] | 0.0064<br>(1.03)<br>[1.01-1.05] | 1.00x10<sup>-8</sup><br>(1.04)<br>[1.03-1.06] | *DLEU1*<sup>g</sup> |
| **rs61975764** | **14:93,014-93,014** | **1/1** | **0.47** | **A/G** | **2.65x10<sup>-8</sup>**<br>**(1.03)**<br>**[1.02-1.04]** | **1.46x10<sup>-7</sup>**<br>**(1.04)**<br>**[1.03-1.06]** | **7.84x10<sup>-7</sup>**<br>**(1.03)**<br>**[1.02-1.04]** | ***RIN3*<sup>g</sup>** |
| rs4381563 | 15:75,275-75,448 | 19/1 | 0.33 | A/T | 7.81x10<sup>-9</sup><br>(0.97)<br>[0.96-0.98] | 0.00027<br>(0.97)<br>[0.96-0.99] | 3.37x10<sup>-9</sup><br>(0.96)<br>[0.95-0.97] | *PPCDC*<sup>g</sup> |
| **rs12956924** | **18:46,451-46,451** | **1/1** | **0.31** | **A/G** | **1.52x10<sup>-8</sup>**<br>**(1.03)**<br>**[1.02-1.05]** | **2.64x10<sup>-5</sup>**<br>**(1.04)**<br>**[1.02-1.05]** | **2.09x10<sup>-7</sup>**<br>**(1.03)**<br>**[1.02-1.05]** | ***SMAD7*<sup>g</sup>** |
| **rs10419921** | **19:16,412-16,412** | **1/1** | **0.30** | **T/C** | **1.67x10<sup>-8</sup>**<br>**(1.03)**<br>**[1.02-1.05]** | **5.62x10<sup>-6</sup>**<br>**(1.04)**<br>**[1.02-1.06]** | **1.09x10<sup>-6</sup>**<br>**(1.03)**<br>**[1.02-1.04]** | ***KLF2*<sup>g</sup>** |
| rs6066184 | 20:45,228-45,716 | 26/2 | 0.26 | G/C | 1.19x10<sup>-13</sup><br>(0.95)<br>[0.94-0.97] | 4.27x10<sup>-6</sup><br>(0.96)<br>[0.94-0.98] | 6.55x10<sup>-16</sup><br>(0.95)<br>[0.93-0.96] | *EYA2*<sup>d</sup> |

Loci found only when analysing all three diseases as one phenotype are marked as bold. More details can be found in S3, S12-S13 and S15 Tables.
<sup>a</sup> Defined as SNPs located <1 Mb apart containing at least one significantly associated genetic variant at $P \leq 3 \times 10^{-8}$ .
<sup>b</sup> Total number of SNPs with $P \leq 3 \times 10^{-8}$ within loci.
<sup>c</sup> Total number of independent associations within the locus, based on conditional analysis (14).
<sup>d</sup> Minor allele frequency.
<sup>e</sup> OR for minor allele.
<sup>f</sup> Lead SNP is in LD ( $R^2 > 0.8$ ) with a missense variant.
<sup>g</sup> Gene(s) closest to the lead SNP.
<sup>h</sup> Lead SNP is in LD ( $R^2 > 0.8$ ) with the lead eQTL SNP. Information on tissue type can be found in S3 Table.

### Annotation of Asthma/hay fever/eczema (combined as a single phenotype) SNPs

For 16 of the 110 asthma and/or hay fever and/or eczema associated loci, the lead SNPs overlapped with a lead SNP for an eQTL (Table 4; S3 and S12 Tables). Among the novel loci, one overlapped with an eQTL for *HIST1H2BD* in whole blood (P=1.11×10^−16^). A decreased level of *HIST1H2BD* seems to increase the risk of this combined phenotype (S3 Table). Probable causal missense variants could be observed at 17 out of 110 significant loci and one of these was observed at one of the novel loci located within *TNFRSF8* and was also identified in the asthma analysis above (S15 Table).

### SNP-based heritability

To quantify the SNP-based heritability for asthma and hay fever/eczema, we used LD score regression analysis (LDSC) (19). These analyses included the same cases and controls as for the association analysis (see Methods). The SNP-based heritability was estimated to be 21% for asthma and 16% for hay fever/eczema (Table 5). Our significant loci, which were located ≥ 1 Mb apart and contained at least one significantly associated genetic variant at P≤3×10'^8^, explained 4.2% of the heritability for asthma and 3.6% of the heritability for hay fever/eczema (Table 5).

**Table 5.** SNP-based heritability in UK Biobank for asthma and hay fever/eczema (combined as one phenotype) estimated with LDSC (19).

|  |  |  | Prevalence used in LD-score<sup>a</sup> |  | (A) All SNPs |  | (B) Without significant loci |  | h2 explained by significant loci |
| --- | --- | --- | --- | --- | --- | --- | --- | --- | --- |
|  |  |  | Population | Sample | h2 | SE | h2 | SE | Absolute terms: (A) - (B) |
| Phenotype | N cases | N controls |  |  |  |  |  |  |  |
| Asthma | 41,926 | 239,751 | 0.117 | 0.117 | 0.210 | 0.017 | 0.168 | 0.010 | 0.042 |
| Hay fever/Eczema | 84,034 | 239,751 | 0.232 | 0.232 | 0.160 | 0.010 | 0.124 | 0.007 | 0.036 |

<sup>a</sup>LDSC requires values for population and sample prevalence when estimating SNP heritability. In these analyses we used the prevalence in the UK Biobank cohort (Table 1) as the sample prevalence. However, due to the cross-sectional design of the UK Biobank we used the same values as population prevalence.

### Identification of phenotype specific loci (SNP)

In our GWA studies, we included all individuals reporting either asthma (for the asthma GWA study) or hay fever/eczema (for the hay fever/eczema GWA study) as cases, independent on if they reported having the other disease phenotype (i.e., asthma cases could have reported having asthma and hay fever/eczema or only asthma). To investigate possible phenotype-specific SNPs, we performed polytomous (multinomial) logistic regression to identify whether the effect of a locus (lead SNP) was significantly (FDR≤0.05) larger for one disease phenotype as compared to another. These effects can therefore be considered as being disease/phenotype specific. To conduct these analyses, we used four non-overlapping groups: 1) asthma cases without hay fever/eczema (N=22,858), 2) hay fever/eczema cases without asthma (N=65,063), 3) asthma cases with hay fever/eczema (only including N=19,299 participants that had reported asthma in combination with hay fever or eczema), and 4) controls without asthma, hay fever and eczema (N=240,817) (Figure 2). Hay fever and eczema were not separated in this analysis due to the small sample size (Table 1). Groups were compared in a pairwise fashion (S16 Table).

A total of 154 lead SNPs (see method section for a description of the selection of lead SNPs), representing the 138 different loci, identified in the GWA study for asthma, hay fever/eczema, or for asthma/hay fever/eczema, were included in polytomous logistic regression analyses. To illustrate the specificity in the Venn diagrams (Figure 3), each SNP was assigned to an area that represents either a phenotype-specific effect (significantly larger in one group of cases) or a shared effect (no significant difference between the two groups of cases).

**Figure 3.**
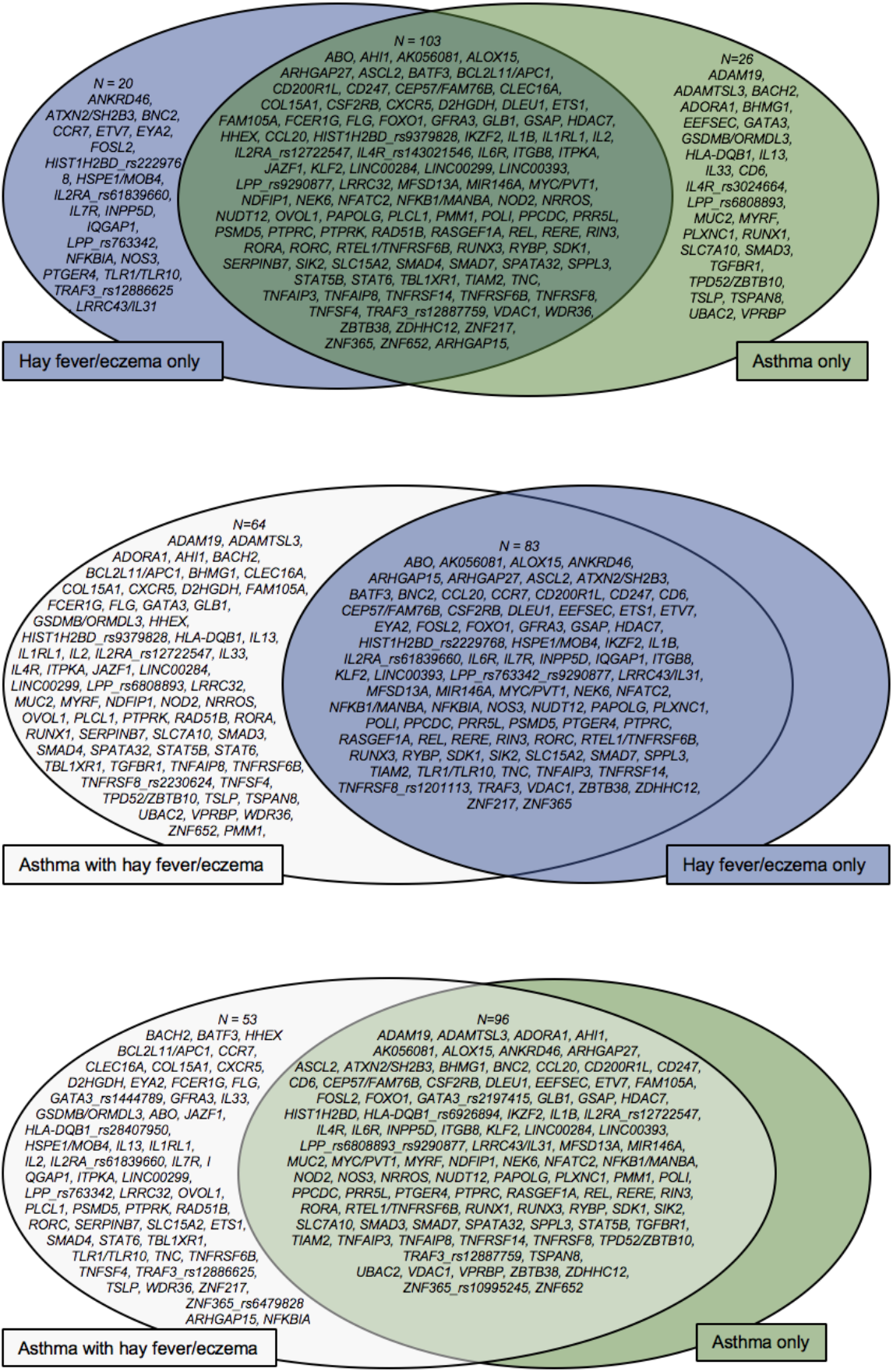
Venn diagram showing the phenotype-specificity of the GWA loci, based on the results from the polytomous logistic regression analyses. The Venn diagram show loci (SNPs) that are specific (significantly larger effect) to or shared between (no significant difference in effects) between two non-overlapping groups of cases. The name of each locus is denoted by the most likely gene(s). At some of the loci (e.g. *IL2RA, LPP*, and *IL4R*), more than one independent (R2<0.8) lead SNP has been analysed in the polytomous logistic regression. If those showed different specificity pattern, they have been included twice in the figure with the name of respective lead SNP(s) also included in the locus name. P-values and estimates for the genes can be found in Table S16 where the area number 1 (green in the figure) indicates specificity for the asthma only, 2 (blue in the figure) specificity for hay fever/eczema only and area number 3 (white in the figure) specificity for asthma with hay fever/eczema (significantly larger estimate in the asthma with hay fever/eczema group of cases).

In the comparison of asthma-only, i.e. without hay fever and eczema, to hay fever/eczema-only, i.e. without asthma, 26 loci/SNPs were specific for asthma-only, i.e. had a significantly higher OR for asthma compared to for hay fever/eczema, while 20 loci/SNPs were specific to hay fever/eczema-only. A major part of the loci/SNPs, 103, showed no significant difference in effect between these disease phenotypes. (Figure 3; S16 Table).

When comparing subjects with asthma and hay fever/eczema to subjects with asthma, 53 loci/SNPs were specific for asthma with hay fever/eczema. No SNP was specific for the asthma only group (Figure 3; S16 Table). For the remaining 96 loci/SNPs, there was no significant difference in effect between subjects with asthma only and subjects with asthma as well as hay fever/eczema.

Finally, when comparing cases of hay fever/eczema only with cases of hay fever/eczema combined with asthma (Figure 3; S16 Table), 64 loci/SNPs had significantly larger effect in the group with hay fever/eczema combined with asthma. No locus had a larger effect in the hay fever/eczema without asthma group. As many as 83 loci/SNPs had no detectable difference in effect between these two disease phenotypes.

For some loci, multiple, possibly independent (R^2^≤0.8) SNPs were included in the analyses. For most of the analyses, such independent SNPs within the same locus showed the same phenotype specificity, or lack of specificity. That is, all independent SNPs within one locus belong to the same area in the Venn diagram (Figure 3). However, for a number of loci, the effect for the different independent SNPs showed different phenotype specificity. This resulted in 149 independent loci/SNPs when comparing the asthma-only group to the hay fever/eczema-only group and when comparing subjects with asthma and hay fever/eczema to subjects with asthma only (Figure 3). For the last group, when comparing cases of hay fever/eczema only with cases of hay fever/eczema combined with asthma, 147 independent loci/SNPs were identified and included in the Venn diagram (Figure 3). For example, two uncorrelated SNPs (R^2^<0.05) were found to be located within the same intron of *IL2RA:* rs61839660, which was associated with hay fever/eczema, and rs12722547, which was associated with asthma in the GWA study. The rs61839660 SNP has a significantly larger effect in both hay fever/eczema-only and hay fever/eczema with asthma, compared to asthma-only but no difference in effect between hay fever/eczema with or without asthma. The effect of rs12722547 was instead significantly larger in the hay fever/eczema with asthma group, compared to the hay fever/eczema without asthma group. Rs12722547 also exhibited a trend (nominal P = 0.05) towards having a larger effect in asthma-only compared to hay fever/eczema-only (S16 Table).

## Discussion

In this large GWA study, including 346,545 unrelated Caucasian participants from UK Biobank, we identified 141 unique loci that are associated with self-reported asthma, hay fever, and/or eczema when these traits are analysed separately or together as combined phenotypes. In comparison with previous studies based on UK Biobank and similar disease phenotypes, our study has several strengths and presents additional results. Out of all identified loci, as many as 41 are novel to our study and have not been reported to be associated with the same disease phenotype previously. Compared to Ferreira *et al* (6) and Zhu *et al* (9), that only included the first release of UK Biobank, we included the full UK Biobank cohort. We also present five different GWA studies for five different phenotypes and further had the strength to identify a number of possible phenotype specific effects that had not been discussed previously.

The largest number of loci were associated with combined phenotype (asthma and/or hay fever and/or eczema), most likely due to the larger sample size of this group. However this is in agreement with a shared genetic contribution between diseases, as has been shown in Ferreira *et al* (6) and Zhu *et al* (9). With this combined phenotype (asthma and/or hay fever and/or eczema), we identified four novel loci that were not found for asthma or hay fever/eczema when analysed separately. Three of these loci appear to be highly relevant to the pathogenies of all three diseases: *SMAD7, KLF2* and *RIN3*. The variant at the *KLF2* locus is located in the 5’ UTR of *KLF2*. This gene plays a role in processes during development including epithelial integrity, inflammation, and T-cell viability. Previous studies have found associations between this locus and lymphocyte percentage of white cells, neutrophil percentage of white cells, white blood cell counts, monocyte percentage of white cells, and eosinophil percentage of granulocytes (20). The variant at the *SMAD7* locus is located within an intron of *SMAD7*. This gene has previously been associated with inflammatory bowel disease (IBD) (21), colorectal cancer (22), and haemoglobin concentration (20). The variant at the *RIN3* locus, is located within an intron of *RIN3*, and is also associated with *RIN3* expression. This gene has previously been associated with myeloid white cell count, eosinophil basophil counts (20), and chronic obstructive pulmonary disease (23). It is worth noting that some of our novel loci have previously been associated with a related phenotype (S1, S5, S8, S10 and S12 Tables). For example, some of the novel asthma loci has previously been associated with IgE levels, eosinophil counts or dermatitis and some of the novel hay fever/eczema loci with IgE levels or eosinophil counts.

For four of the novel loci: near *TNFRSF8, MYRF, TSPAN8*, and *BHMG1*; the lead SNP was in LD with potentially deleterious missense variants. The lead genetic variant at the *TNFRSF8* locus, rs2230624, which is associated with asthma as well as the combined asthma/hay fever/eczema phenotype, is a potentially causal missense variant that causes a cysteine to a tyrosine substitution in the TNFRSF8 protein. This protein, which is also referred to as *CD30*, is a receptor that is expressed on activated T and B cells and has been shown in clinical studies to have a role in the development of allergic asthma (24). To the best of our knowledge, this is the first time that this locus has been associated with asthma and allergy in a GWA study. The lead SNP at the *MYRF* asthma locus, rs174535, is a missense variant within the myelin regulatory factor protein (*MYRF*) that causes a serine to arginine substitution near the end of the protein. This gene lies within the fatty acid desaturase (*FADS*) cluster on a fatty acid synthesis-associated haplotype (25). Variants on this haplotype are also strongly associated with expression of *FADS1* and *FADS2*, two genes that are involved in the desaturation of polyunsaturated fatty acids in the biosynthesis of long chain polyunsaturated fatty acids (LC-PUFAs). One of these variants has previously been shown to modulate the effect of breast-feeding on asthma (26); another has been associated with increased risk of inflammation (27). Reduced capacity to desaturase omega-6 LC-PUFAs due to *FADS* polymorphisms has been shown to be nominally associated with reduced risk for development of eczema, potentially due to a pathogenic role of omega-6 LC-PUFAs in development of allergy (28). For seven of the 41 novel GWA loci, the lead SNP was in LD (> 0.8) with an eQTL. We could see a positive correlation between expression of *TUBG2* (in skin), *HHEX* (in cell transformed fibroblasts), *EEFSEC* (in lung and cell transformed fibroblasts), and *ADAM19* (in whole blood) and risk of disease, as well as a negative correlation for *TMEM258* and *HIST1H2BD*. Decreased expression of *TMEM258* in cell transformed fibroblasts was associated with increased risk of asthma. In transgenic experiments in mice, it has been shown that a lower expression of *TMEM258* leads to severe intestinal inflammation (29), which agrees with our results. A possible limitation of this analysis is that it relied solely on the GTEx database. Additional sources of information on eQTLs may increase the total number of eQTLs that are associated with asthma, hay fever and eczema.

For 16 loci that were associated with asthma, 20 loci associated with hay fever/eczema and for 21 loci associated with asthma/hay fever/eczema, we identified multiple independently associated variants. This indicates that several of the asthma-, hay fever-, and eczema-associated loci represent multiple independent disease-associated variants. As an example, the *FLG* locus contains three independent asthma, and asthma/hay fever/eczema-associated variants. This gene has previously been shown to contain loss-of-function mutations that are causal for skin barrier deficiency and strongly predispose to both eczema and asthma (30). The four most prevalent European *FLG* mutations are c.2282del4, p.R501X, p.R2447X, and p.S3247X (30). An additional example is the *HLA* region whose association with immune diseases is particularly complex and which has previously been suggested to include several independent regulatory factors (31). In our analyses, we identify as many as 21 independent associations within this locus.

As highlighted by this study, as well as previous studies (2–7, 32), many disease-associated loci overlap between asthma, hay fever and eczema. However, several loci were only significantly associated with only one of the investigated phenotypes. By testing for association with hay fever and eczema separately in a smaller set of participants, we were able to resolve some of these signals. Interestingly, one of the strongest associations for hay fever/eczema (P=7.96×10^−25^), found within the *FLG* locus, was more significantly associated with eczema when this phenotype was analysed separately (P=1.15×10^−69^). In contrast, this variant was not associated with hay fever when hay fever was analysed separately. It was, however, associated with asthma (P=2.37×10^−27^). This is in agreement with the previous GWA study by Ferreira *et al*, where a SNP at the *FLG* locus was shown to be specifically associated with eczema (6). However, a different study has shown that mutations within the *FLG* locus are associated with eczema starting in the first year of life, and that these mutations are associated with a later development of both asthma and hay fever (33). This is an example of the typical progression of allergic diseases that often begin early in life, which is commonly referred to as the atopic march (33–35). When analysing hay fever separately, we identified one novel locus near *CBLN1*. Studies on transgenic mice have shown that knock-out of *CBLN1* mimics loss-of-function mutations that occur in the orphan glutamate receptor, *GRID2* (36). Autoantibodies against glutamate receptors are involved in the development of autoimmune disease (37). One novel locus was also identified and replicated when analysing eczema separately, downstream of *TAGAP*. This locus has previously been associated with celiac disease (38) and multiple sclerosis (39).

We further investigated our novel asthma, hay fever/eczema and eczema loci in two independent cohorts. We were only able to replicate six SNPs out of the 15 identified to be novel for asthma with the summary statistics from the GABRIEL asthma consortium. The GABRIEL GWA study only included 582, 802 SNPs genotyped with the Illumina Human610 quad array, and therefore, a large number of SNPs did not overlap between our studies. However, for most of the loci where we identified the same SNP or a proxy in LD (R2≥0.8), we did find a nominal replication (P≤0.05) (Table 2). The GABRIEL study was based on childhood asthma while the UK Biobank asthma is based on adult asthma and these two disease phenotypes may therefore have some different underlying genetic effects. It is also important to remember that a lack in replication may also be due to a lower power to detect associated SNPs in GABRIEL due to a smaller sample size. Five lead SNPs identified for the combined analysis hay fever/eczema replicated in the EAGLE study (Table 3). The lack of replication is most probably due to differences between phenotypes. While the EAGLE study only included eczema cases, our study also included hay fever. All SNPs that did not replicate in EAGLE, was neither statistically significant when analysing eczema independently in UK Biobank (Table 3). We also tried to replicate the four novel loci that was only identified in the combined analysis, asthma/hay fever/eczema, using the GABRIEL and EAGLE cohorts. However, none were nominally replicated (P‹0.05) which is probably due to the smaller sample sizes in GABRIEL and EAGLE, and most importantly due to the differences in disease phenotypes between the cohort.

Out of all asthma and/or allergic disease-associated loci that have been reported to the GWAS catalog as of December 2, 2018, the majority (N=108) were nominally replicated in our study (P≤0.05; S14 Table). Twelve associations were not possible to test due to lack of data, i.e. neither the reported SNP nor any SNP in LD with the reported SNP were presented in our data (S14 Table). Asthma, hay fever, and eczema are known to be heterogeneous diseases in which environmental factors play an important role (1). Genetic variants associated with asthma, hay fever and eczema are likely to be population specific (40). It is therefore possible that population-specific variants are not detected in our study. Many of the previous associations that were not replicated in our study have been identified in studies that have used a somewhat different phenotype (41, 42), populations of different ancestry (15, 43) or small sample sizes (< 10,000) (43, 44). Research findings from studies on smaller cohorts are more likely to be false positives, especially when no replication of primary findings has been performed, and are thereby less likely to represent true causative mechanisms (45) (for more information, see S14 Table). A recent GWA study by Zhu *et al* (9), which was also conducted on the UK Biobank cohort, however using a different combination of allergies as a phenotype, reported seven novel allergy-associated loci, five of which were replicated in our study. These loci where not available in the GWAS catalog at the time of writing this article and are therefore not included in S14 Table. The two loci that did not replicate in our study where mapped to *ALG9* on chromosome 11(rs659529) and to *EVI5* on chromosome 1 (rs12743520).

In previous GWA studies for asthma, the disease phenotype commonly contained other disease phenotypes as well, e.g. participants with asthma commonly also report hay fever/eczema. In contrast, our polytomous logistic regression approach allowed for identification of genetic variants with differing effects between the different sub-phenotypes. These effects can therefore be considered as being disease/phenotype specific. This was achieved by subdividing the participants in four non-overlapping groups depending on asthma and hay fever/eczema status. The SNPs that were included in these analyses were selected from our main GWA analyses, but not including the two SNPs identified for the hay fever and eczema phenotypes analysed separately since we did not have power (large enough sample size) enough to include hay fever and eczema separately in these analyse. This means that a locus that was defined as specific for asthma-only has already been associated with any of the combined phenotypes and/or with asthma, independent of hay fever /eczema status. The association for such variants may have been due to comorbidity between asthma and the other diseases, e.g. a larger fraction of asthma cases in the hay fever/eczema group compared to the controls, or that the effect of the asthma-only specific variants was only partly diluted by being combined with other disease phenotypes. A large number of loci exhibited differential effects between hay fever/eczema-only and asthma-only. As many as 20 loci had a significantly larger effect on hay fever/eczema-only compared to their effects on asthma while 26 loci exhibited larger effects on asthma compared with their effects on hay fever/eczema (Figure 3). Among the loci that were specific for asthma-only, we find *ADAM19* and *ADAMTSL3* which are proteins with multiple biological roles within the cell and believed to be important in a number of diseases, including asthma (46). Among the loci that were specific for hay fever/eczema we find the toll like receptor loci, *TLR1/TLR10*, which also showed a larger effect on hay fever compared to asthma-only in the Ferreira *et al* study (6). Most associated variants at this locus are located within the promoter region of *TLR1*, which encodes the toll-like receptor 1. This protein constitutes a component of the innate immune response to microbial pathogens (47). Several loci that overlap between asthma-only and hay fever/eczema-only were annotated to genes related to tumor necrosis factor (TNF) function, such as *TNFAIP3, TNFAIP8, TNFRSF11A, TNFRSF14, TNFRSF6B, TNFRSF8, TNFSF4*. These proteins are mainly expressed in immune cells and regulate immune response and inflammation as well as proliferation, apoptosis and embryogenesis (48).

The largest number of phenotype-specific loci was observed for the group of cases with asthma and hay fever/eczema (Figure 3; S16 Table), a group of cases that has not been included in similar analyses in previous studies (6). This is a group of participants with an allergic disease in combination with asthma, which could to some degree represent participants with allergic asthma. The number of phenotype-specific loci is considerable larger in our study compared to previous studies that have performed similar analyses, such as the study by Ferreira *et al*. (6),which only identified six disease specific loci. This is not surprising since our analyses included larger sample sizes: N= 65,063, N=22,858, and N=19,299 compared to N= 33,305, N=12,268, and N=6,276 in the study by Ferreira *et al* (6) for the three sub-groups included in the analyses of disease-specific effects. In addition, since only genome-wide significant SNPs were taken forward to the polytomous logistic regression analyses, we used the False Discovery Rate by Benjamini-Hochberg to adjust for multiple testing. This increases the power to pinpoint as many positive findings as possible, still with a small false-discovery rate (5% in our case), compared to the more conservative Bonferroni method used in the previous study by Ferreira *et al* (6). The previous study, also separated hay fever and eczema, and compared the three groups hay fever-only, eczema-only and asthma-only. Since different subgroups of cases were analysed in our study our results do not disagree with that of Ferreira *et al* (6) that found six disease-specific SNPs: near *FLG, RPTN-HRNR* (close to *FLG), IL2RA, IL1RL2-IL8R1, WDR36-CAMK4* and *GSDMB;* where five of them were significantly different between hay fever and eczema.

The SNP-based heritability was estimated to be 21% for asthma and 16% for hay fever/eczema. These percentages represent the portion of heritability that can be captured by the common genetic variants that were included in the GWA study. The SNP-based heritability for asthma has previously been estimated at 15%, which is slightly lower than the estimate from our study and probably due to a smaller sample size, and/or a difference in disease definition in the previous study (6). In comparison to the high estimates for the heritability (33-95%) from family and twin studies (1, 10, 12), this suggests that a major contribution to the genetic risk for asthma, hay fever and eczema might not be identified in studies using common genetic variants, or need cohorts with even larger sample sizes. However, heritability estimates from family and twin studies have been suggested to be overestimated (49–51) due to the fact that these estimates often are based on simplistic models that ignore shared environmental factors. Our estimate might also be lower due to the presence of disease-associated rare variants that are not captured by the SNP-based heritability estimate.

A possible limitation of the present study is the self-reported phenotypes, which might lead to a recall bias and misclassification. Another limitation is that the UK Biobank cohort traits are not independent since there are shared cases between asthma, hay fever and eczema and completely shared controls. However, findings presented in this article apply to a single large population of individuals of similar age. Population stratification was also controlled for by filtering for Caucasian participants, including ancestry derived principal components and adjusting for the LD-score intercept in our analyses. Participants of the UK Biobank are also more likely to be exposed to more similar environmental factors, compared to the participants of previous meta-analyses that utilise a large number of smaller cohorts from different countries and age-groups. Analysing hay fever and eczema as a combined phenotype is another limitation in our study, which prohibits identification of hay fever- and eczema-specific SNPs. We therefore refer to SNPs as phenotype-, rather than disease-specific in the polytomous logistic regression analyses. However, both hay fever and eczema are IgE mediated hypersensitivities and therefore probably share similar physiology (52).

## Conclusion

In summary, we describe 15 novel loci for asthma, 22 novel loci for hay fever and/or eczema and an additional four novel loci were found when analysing asthma, hay fever and eczema together. Two novel loci were also identified when analysing hay fever and eczema separately. Pinpointing candidate genes for common diseases are important for tailor-made studies that want to prioritize candidate genes for developing novel therapeutic strategies. This study further highlights a large amount of shared genetic contribution to these diseases, indicating that the comorbidity between asthma, hay fever and eczema is partly due to shared genetic factors. However, we also show that a number of SNPs have a significantly larger effect on one of the phenotypes, suggesting that part of the genetic contribution is phenotype specific.

## Methods

### Study population

The UK Biobank includes 502,682 participants recruited from all across the UK. Participants were between 37 and 73 years old at time of recruitment between 2006 and 2010. Most participants visited the centre once, but some individuals visited the centre at up to three times. Participants answered questions about self-reported medical conditions, diet, and lifestyle factors. A total of 820,967 genotyped SNPs and up to 90 million imputed variants is available for most participants. The UK Biobank study was approved by the National Research Ethics Committee (REC reference 11/NW/0382). An application for using data from UK Biobank has been approved (application nr: 15479). We included 346,545 unrelated Caucasians (see selection of participants and sample QC below) with genotypes from the second UK Biobank genotype release (Table 1).

### Disease phenotypes: asthma and hay fever/eczema

Self-reported asthma as well as self-reported hay fever and/or eczema (combined) were assessed using the UK Biobank touch screen question number (Data field 6152), which asked the participants the following question: *has a doctor ever told you that you have had any of the following conditions? (You can select more than one answer):* 1) asthma and 2) hay fever, allergic rhinitis or eczema, 3) none of the above or 4), prefer not to answer. Because hay fever and eczema diagnosis could not be separated we called this variable hay fever/eczema (i.e., participants reported hay fever and/or eczema). All participants were also invited to participate in an interview. At first, nurses (trained UK Biobank staff-member) confirmed with each participant that the information they provided on the screen or questionnaire was correct if they had answered that a doctor had told them they had one or more of the following diseases: heart attack, angina, stroke, high blood pressure, blood clot in leg, blood clot in lung, emphysema/chronic bronchitis, asthma, or diabetes. Due to the confirmation of asthma cases, the overlap in asthma variables between the touch-screen questionnaire and verbal interview was very high. For asthma, only 622 individuals were removed due to conflicting answers between the touch-screen and verbal interview. Using a drop-down menu, the nurses could also add other diagnoses. These diagnoses (UK Biobank data field 20002) were used to define hay fever and eczema cases separately. However, the disease prevalence in this variable appears to be largely underreported as many individuals reported hay fever or eczema in the touch-screen questionnaire but did not report hay fever or eczema during the interview. For this reason, the touch-screen data variables hay fever/eczema, with a much larger sample size (Table 1) compared to hay fever and eczema separately, was used as one of the primary phenotypes analysed in this study. For hay fever/eczema, 4,881 individuals were removed due to conflicting answers between the touch-screen questionnaire and the interview, for individuals reported they had hay fever during the interview but not on the touch-screen questionnaire (N=2,143), or reported they had eczema in the interview but not in the touch-screen (N=2,738). We further removed 22 individuals who had asked to be removed from the UK Biobank.

### Controls

Controls (N=239,773) were selected as individuals answering “none of the above” in question 6152, and who did not report asthma, hay fever or eczema in variable number 20002. The same controls were used for all phenotypes.

### Genotyping

The UK Biobank Axiom array had been used to genotype 438,417 of the 502,682 UK Biobank participants. The other 49,994 samples (all from the interim release) had been genotyped on the closely related UK BiLEVE array. The UK BiLEVE cohort and the rest of the UK Biobank differ only in small details of the DNA processing stage. The two arrays have 95% common marker content. We included a variable for array type (UK BiLEVE or UK Biobank Axiom) as covariate. SNPs in UK Biobank were imputed using UK10K (53) and 1000 genomes phase 3 (54) as reference panels. Imputation in the second release resulted in 92,693,895 SNPS (released in June 2017). However, because the UK Biobank reported problems with imputation quality for a subset of the SNPs (caused by mismatch in coordinates between the UK10 and the 1000 genomes reference panels), we followed the recommendation to only include genetic variants included on the HRC panel (55) (N=39,727,058).

### Quality control

Quality control of genotype data and imputation of genotypes had already been carried out centrally by UK Biobank. From the imputed dataset, we only included SNPs in the HRC panel with a MAF ≥ 0.01. We removed SNPs deviating from Hardy-Weinberg (P-value < 1×10^−20^) and markers with more than 5% missing genotype data. We only included SNPs with an imputation quality >0.3. After QC, a total of 15,688,218 autosomal SNPs and SNPs on the X-chromosome were included in our analyses. We only included Caucasian participants who were clustering according to the genetic principal components (56,180 non-Caucasians were removed: individuals listed in UK Biobank data file 22006). We further removed first and second-degree relatives (N=32,751), using kinship data (estimated genetic relationship > 0.044), and participants with sex discordance, high heterozygosity/missingness (individuals listed in UK Biobank data field 22010 and 22027), and participants with more than 5% missing genotypes. After QC and exclusion, 346,545 unrelated Caucasian participants remained.

### Genome-wide association study

A GWA study were performed for each phenotype using logistic regression and an additive genetic model implemented in PLINK version 1.90 (56). We performed a GWA studies for five sets of phenotypes: 1) asthma (independent on hay fever and eczema status), 2) hay fever/eczema (hay fever and/or eczema independent on asthma status), 3) hay fever and/or eczema and/or asthma, as well as 4) hay fever (independent on asthma and eczema status) and 5) eczema (independent on asthma and hay fever status). The same controls, that have reported that they did not have any of the disease phenotypes, were used for all analyses (N=239,773). The following covariates were included in our analysis: Townsend deprivation index (TDI) (as a proxy for socioeconomic status), sex, age, smoking, and the first ten ancestry derived principal components. In addition, to adjust for the different genotyping chips, we included a binary indicator variable for UK Biobank Axiom versus UK BiLEVE genotyping array. We calculated the LD-score intercept, using the LD score regression software (LDSC) (19), for each phenotype and adjusted the summary statistics accordingly (19). The genome-wide significance threshold was set to 3×10^−8^, as suggested for GWA studies that include variants with a minor allele frequency (MAF) ≥ 0.01(57), which was the threshold used in our study. Individual loci were defined as regions with at least one significantly associated SNP (P ≤ 3×10^−8^). Start and stop positions for each locus were where no additional significantly associated SNPs could be found (upstream for start position, or downstream for stop position) within 1 Mb.

### Identification of additional independent variants within associated loci

To identify independently associated variants within each defined locus (significant SNPs (P ≤ 3×10^−8^), within 1 Mb), we used an approximate conditional analyses implemented in GCTA (14). LD calculations were based on 5,000 randomly selected Caucasian participants from UK Biobank (after sample QC). For each locus, the most significant top SNP was identified and the summary statistics of all SNPs within the same locus was adjusted by the effect of the lead SNP. After adjusting for the lead SNP, we identified the most significantly associated SNP within the locus that remained significant (P ≤ 3×10^−8^). In the next step, we once again adjusted the summary statistics of all SNPs within the same locus, by including the effect of both the original lead SNP and the conditional lead SNP form the first iteration. This process was thereafter repeated until no other SNPs within the locus were found significant after adjusting for all previously detected independent lead SNPs.

### Determining the novelty status of significant loci

To determine whether significant loci were novel to any of the diseases, we compiled a list of all asthma, hay fever, eczema and allergy risk SNPs with genome-wide significant association (≤ 5×10^−8^) reported in the NHGRI-EBI GWAS catalog (downloaded December 2, 2018). We also searched for GWA study results using PubMed and bioRxiv. We classified a locus to be ‘novel, if the locus was > 1 Mb from any of the previously reported loci/variants for the disease. We also estimated LD between each lead SNP and all genome-wide significant associations found in the NHGRI-EBI GWAS catalog, to determine whether the lead SNP was a novel variant in a known locus (if the locus was < 1 Mb from any of the previously reported loci/variants for the disease). We considered our associated SNP to be a novel variant if R^2^ were smaller than 0.05 between our top associated variant and previously reported variants within the same locus. A locus was also reported as novel for a specific disease (i.e., asthma) if previous GWA studies only reported association to a different allergic disease (for example hay fever). If the locus was previously reported for a combined phenotype, i.e. in studies combining different allergic diseases, including the one tested, it was not reported as a novel locus.

### Annotation of target genes and identification of causal genetic effects

To identify likely target genes for associated variants, we first reported the closest gene(s) to the lead SNP for each locus and reported if the SNP was intronic or exonic using the Human Genome Browser (GRCh37). We also performed additional analyses to potentially better define plausible target genes. To examine the relationship between the lead SNP for each locus and gene expression we used the Genotype-Tissue Expression (GTEx) database (58) to find evidence of overlap with expression quantitative loci (eQTLs). We downloaded significant eQTLs from the Genotype-Tissue Expression (GTEx) database. First, we selected GTEx SNPs that overlapped with the UK Biobank SNPs and used a conservative significance threshold P ≤ 2.3 × 10^−9^ for cis effects (<1 Mb) form the GTEx data, in agreement with previous studies (6). Second, we identified the most significant eQTL SNP for each tissue and gene in the GTEx dataset. Third, we estimated the LD between the lead eQTL SNPs and our lead GWA study SNPs. A lead GWA SNP in LD (R^2^ > 0.8) with a lead GTEx eQTL SNP was considered to overlap with the eQTL. Only cells or tissues that were relevant for our disease phenotypes were considered when searching for eQLTs, including EBV-transformed lymphocyte, transformed fibroblasts, whole blood, lung and skin (sun exposed and not sun exposed).

We also used the Bioconductor biomaRt (59) package in R for functional annotation of associated SNPs. In BiomaRt, lead SNPs, and all SNPs in LD (R^2^>0.8) with a lead SNP, were cross-referenced against: Ensembl Genes, Ensembl Variation, and Ensembl Regulation version 91 (Accessed 9 December 2017 using the human assembly GRCh37). Here we checked whether the lead SNPs were in LD (R^2^>0.8) with a potentially functional genetic variant by investigated regulatory features for the SNPs (i.e., promoters, enhancers etc.), binding motifs (i.e., if any of the SNPs were found within a motif for a transcription factor), and if the SNPs were possibly damaging variants (i.e., missense, stop gained, stop lost, or splice acceptor/donor variants) and if the variants were predicted to be deleterious by SIFT or PolyPhen.

### Replication

We replicated our novel asthma, hay fever/eczema and eczema loci in two independent cohorts the EAGLE eczema consortium and the GABRIEL asthma consortium (P≤0.05). Our novel loci identified for asthma was replicated using the summary statistics from the GABRIEL consortium which consisted of 10,365 physician-diagnosed asthmatic cases and 16,100 healthy controls (60). All individuals in GABRIEL were genotyped for 582,892 SNPs using the Illumina Human610 quad array. More information on this cohort has been published elsewhere (60). The EAGLE consortium GWA summary statistics consists of 21,000 atopic dermatitis (eczema) cases and 96,000 controls (61) and were used to replicate novel loci for hay fever/eczema and eczema analysed separately. Further information about this cohort has been published previously (61). If the lead SNP from our study was not found in GABRIEL or EAGLE, we search for a proxy in LD (≥0.8) with the lead SNP.

### SNP-based heritability

To quantify the SNP-based heritability for asthma and for hay fever/eczema (combined as a single phenotype) we used LD score regression software (LDSC) (19) including the same cases and controls as for the association analysis for each phenotype (19). To calculate the heritability on the liability scale, we needed to adjust for disease prevalence. Since this was a population-based study, we set the Caucasian population and sample prevalence to the one calculated for each disease in UK Biobank. We included 1,108,908 HapMap SNPs to calculate the heritability for asthma and hay fever/eczema. We also removed all significant loci from each individual GWA study result to estimate how much of the heritability was explained by the significant loci reported in this study.

### Identification of phenotype-specific loci (SNP)

To identify possible phenotype-specific SNPs, we performed polytomous (multinomial) logistic regression to identify whether the effect of a locus (lead SNP) was significantly (FDR≤0.05) larger for one disease phenotype as compared to another. These effects can therefore be considered as being disease/phenotype specific. To conduct these analyses, we used four non-overlapping groups: 1) asthma cases without hay fever/eczema (N=22,858), 2) hay fever/eczema cases without asthma (N=65,063), 3) asthma cases with hay fever/eczema (only including N=19,299 participants that had reported asthma in combination with hay fever or eczema), and 4) controls without asthma, hay fever and eczema (N=240,817) (Figure 2). Hay fever and eczema were not separated in this analysis due to the small sample size (Table 1).

We performed polytomous logistic regression for all possibly independent (R^2^<=0.8) associated lead SNPs identified in the asthma, hay fever/eczema or asthma/hay fever/eczema GWA studies. For some regions, different SNPs, that represent the same signal (R^2^>0.8 between the SNPs), were identified in the different GWA studies. For these regions, only the SNP with the lowest P-value from the original GWA study was included in these analyses. For regions where, different lead SNPs were identified in the different GWA studies, and where these lead SNPs were not in strong LD (R^2^<=0.8), all lead SNPs were included in the analyses.

The polytomous (multinomial) logistic regression was performed with the response variable, *Y*, being categorically distributed with K=4 non-overlapping groups/outcomes (the four non-overlapping groups are explained above). Out of K·(K-1)/2=6 comparisons in total, there are K-1=3 independent comparisons. The logit function is defined as the logarithm of the quotient between the probability of a given outcome (e.g., P(Y=1)) and the probability of a reference or pivot outcome (i.e., P(Y=4) in our case). This function is assumed to be linear in all explanatory variables, including covariates and the specific SNP under consideration. Note that the beta estimates (i.e., the log-odds ratios) are unique for each comparison. The polytomous (multinomial) regression was performed using multinom in the R library nnet for the three independent odds: P(*Y*=1)/P(*Y*=4), P(*Y*=2)/P(*Y*=4), and P(*Y*=3)/P(*Y*=4). Beta estimates, standard errors, and p-values (two-sided, normal approximation) for the remaining comparisons between phenotypic outcomes (i.e., P(*Y*=1)/P(*Y*=2), P(*Y*=1)/P(*Y*=3), and P(*Y*=2)/P(*Y*=3)) were calculated from the model output such that, e.g., *beta*_12_ = *beta*_14_ – *beta*_24_ and *se*_12_^2^ = *se*_14_^2^ + *se*_24_^2^, where the first subscript denotes the outcome of interest while the second subscript denotes the reference outcome.

To determine whether the lead SNPs were specific to one disease phenotype or shared among phenotypes, we identified for which disease phenotype the OR was the highest (we used the value of the OR rather than the most significant P-value in order not to be influenced by the different power in the phenotype groups due to different sample-sizes), and whether the OR was significantly (FDR ≤ 0.05) higher compared to the other disease phenotypes. As a threshold for significance, we used an FDR (Benjamini-Hochberg) value of 0.05, corresponding to a nominal P-value of < 0.017 in the three sets of cases vs cases analyses. In our analyses, an FDR adjustment is to prefer (in favour of Bonferroni) due to its power to pinpoint as many positive findings as possible, while retaining a low false-discovery rate (5% in our case). Results were plotted as Venn diagrams to show the pair-wise overlap between disease phenotypes. If two SNPs from the same locus that were not in LD with each other (R^2^ <= 0.8) were assigned to the same area, the locus only occurs once in the Venn diagram. However, for a few loci, multiple unlinked (R^2^ <= 0.8) SNPs from the same locus were assigned to different areas. Such loci were included at multiple locations in the Venn diagram together with the name of the SNP (i.e., gene_SNP).

## Ethics

UK Biobank was given ethical approval by the North West Multicentre Research Ethics Committee, the National Information Governance Board for Health and Social Care and the Community Health Index Advisory Group. UK Biobank holds a generic Research Tissue Bank approval granted by the National Research Ethics Service (http://www.hra.nhs.uk/) that lets applicants conduct research on UK Biobank data without obtaining ethical approvals for each separate project. Access to UK Biobank genetic and phenotypic data was given through the UK Biobank Resource under Application Number 15479. All participants provided signed consent to participate in UK Biobank.

## Data availability

The genotypes and phenotypes included in the current study are available from the UK Biobank data, which can be accessed by researchers upon application (https://www.ukBiobank.ac.uk/). Summary statistics and codes used for this project can be accessed by contacting the corresponding author.

## URLs

UK Biobank, http://www.ukBiobank.ac.uk; PLINK, https://www.cog-genomics.org/plink2; NHGRI-EBI GWAS Catalog, https://www.ebi.ac.uk/gwas/; Software tool for LD Score estimation and estimation of variance components from summary statistics, https://github.com/bulik/ldsc/; GCTA, http://cnsgenomics.com/software/gcta/; BiomaRt, http://www.bioconductor.org; GTEx, https://www.gtexportal.org/home/; DGIdb, http://www.dgidb.org/.

## Acknowledgement

We acknowledge all the participants and the administrative staff at the UK Biobank. The computations were performed on resources provided by SNIC through Uppsala Multidisciplinary Centre for Advanced Computational Science (UPPMAX) under projects b2016021, b2017059, sens2017538, and sens2017541. The work was supported by grants from the Swedish Society for Medical Research (SSMF), the Kjell and Märta Beijers Foundation, Göran Gustafssons Foundation, the Swedish Medical Research Council (Project Number 2015-03327), the Marcus Borgström Foundation, the Åke Wiberg Foundation, the Borgström Hedström Foundation and the Swedish Heart-Lung Foundation.

## Author contributions

Planned the study (WEE and ÅJ), analysed the data (WEE, ÅJ, TK), literature search (WEE), Figures (WEE, MRA, ÅJ), data interpretation (WEE, ÅJ, TK, MRA), writing of manuscript (WEE, ÅJ, TK, MRA).

---

**S1 Figure** QQ-plot for Asthma in UK Biobank. The red line denotes the expected null-line of no association.

**S2 Figure** Regional plot for the missense variant rs2230624 in asthma.

**S3 Figure** QQ-plot for Hay fever and/or Eczema (combined) in UK Biobank. The red line denotes the expected null-line of no association.

**S4 Figure** Manhattan plots for Hay Fever and/or Eczema (combined), Hay Fever (only) and Eczema (only) analysed in UK Biobank for autosomal chromosomes. The black horizontal line indicates the genome wide threshold (3×10^−8^). The black regions in the hay fever and/or eczema (combined) plot represent novel loci found in this study and the black regions in the hay fever (only) and eczema (only) plot represent two novel loci, not found in previous GWA studies or in our combined hay fever / eczema analysis.

**S5 Figure** QQ-plot for Hay fever (only) in UK Biobank. The red line denotes the expected null-line of no association.

**S6 Figure** QQ-plot for Eczema (only) in UK Biobank. The red line denotes the expected null-line of no association.

**S7 Figure** QQ-plot for Asthma and/or Hay fever and/or Eczema (combined) in UK Biobank. The red line denotes the expected null-line of no association.

## Abbreviations

SNP: Single nucleotide polymorphism
GWA study: Genome Wide Association Study

